# PlumageParts: A fine-grained avian plumage segmentation dataset and benchmark for ecological image analysis

**DOI:** 10.64898/2026.07.30.741737

**Authors:** Yichen He, Eleftherios Ioannou, Kathryn Harris, Gavin Thomas, Steve Maddock, Julien P. Renoult, Christopher Cooney

## Abstract

Fine-grained localisation of plumage regions is a prerequisite for computational analyses of avian colouration, patterning and visual traits in ecological and evolutionary research. Progress is limited by the scarcity of image resources with annotations aligned to biologically meaningful anatomical units: existing avian benchmarks provide either landmark points or coarse part categories that do not capture ornithologically defined plumage regions. We present a curated dataset of 4,705 bird images annotated for nine plumage regions: head, throat, breast, belly, vent, back, coverts, remiges and tail. Spanning 39 avian orders and 222 families, the dataset provides a taxonomically broad resource for fine-grained avian image analysis. The dataset was built through an iterative model-assisted annotation workflow, in which model predictions were reviewed and corrected rather than drawn from scratch, improving the efficiency of region-level annotation. We benchmark classical segmentation architectures, SAM-based models and self-supervised foundation-model encoders on this task. A frozen DINOv3 encoder with a lightweight decoder achieved the highest performance on the held-out test set, reaching 84.01% mean Intersection over Union while requiring substantially less memory than end-to-end fine-tuning. The model generalised to external avian benchmarks, including the bird subset of PartImageNet and CUB-200-2011, and achieved competitive performance on the full PartImageNet part-segmentation benchmark, which includes diverse animal taxa. We provide a modular detect-track-segment pipeline as a proof-of-concept extension to video data. Together, these results show that anatomically grounded avian annotations can serve both as a resource for plumage phenotyping and as a benchmark for efficient, transferable biological part segmentation.

**Author Summary:** Birds vary enormously in colour and pattern, but studying this variation at large scales requires more than identifying the bird in a photograph. Researchers often need to know where each colour or pattern occurs on the body, such as on the head, throat, breast, wing or tail. We created PlumageParts to make this kind of region-level analysis easier. The dataset contains 4,705 bird images annotated into nine biologically meaningful plumage regions, covering a wide range of bird families and orders. To build the dataset efficiently, we used a model-assisted workflow in which computer-generated masks were checked and corrected by researchers rather than drawn entirely by hand. We then tested several image-segmentation approaches and found that a frozen self-supervised vision model, combined with a lightweight decoder, provided accurate plumage-region predictions while requiring relatively modest computing resources. The same approach also performed well on a broader animal part-segmentation benchmark, suggesting that it may be useful beyond birds when suitable annotations are available. By releasing the annotations, code and trained model, we aim to support future studies of bird plumage and biologically meaningful image segmentation.

## Introduction

Region-level phenotypic analysis is increasingly important for addressing questions in ecology and evolution (Pérez-Rodríguez et al., 2017). Compared with whole-organism metrics (e.g. body size), measurements of specific anatomical components, such as colour patches, patterned areas or defined body regions, can reveal variation across species, sexes or environments that would otherwise be obscured (Cooney et al., 2019; López-Idiáquez et al., 2022). Such information is central to studies of morphological integration, modularity and functional trade-offs among phenotypic components, which can shape evolutionary trajectories under selection (Klingenberg, 2008; Kühl C Burghardt, 2013). To enable these analyses at scale, however, homologous regions must first be localised consistently across large image collections before colour, pattern or shape features can be extracted, a step that remains a major bottleneck in image-based phenotyping (Lürig et al., 2021).

Birds provide a clear example of this challenge. The avian radiation represents an important model system in ecology and evolution, and plumage colouration and patterning specifically are closely linked to selection, signalling, environmental adaptation and species diversification (Hill & McGraw, 2006). Existing broad-scale studies of avian plumage colouration have generated important insights using whole-organism colour metrics that can be automated across large datasets, such as overall brightness or colourfulness (Cooney et al., 2022; Dale et al., 2015; Y. He et al., 2022). In contrast, analyses of specific plumage patches, such as the crown, breast or belly, often require manual delineation from photographs or targeted spectrometry of specific regions. These patch-level data can address more fine-grained questions about sexual selection, signalling function and local adaptation that are not accessible from whole-organism summaries alone (Cooney et al., 2019; López-Idiáquez et al., 2022; Nolazco et al., 2023), but they remain difficult to scale because corresponding anatomical regions must be identified consistently across many individuals and species. Here, we focus on this upstream segmentation problem: assigning pixels to biologically meaningful plumage regions as a prerequisite for downstream colour, pattern or shape analyses.

Image segmentation offers a route to automating this region-localisation step. In computer vision, semantic segmentation assigns a category to every pixel in an image (Long et al., 2015), whereas instance segmentation distinguishes individual object instances of the same class (K. He et al., 2017). Our work focuses on semantic part segmentation (J. He et al., 2023): rather than separating multiple bird individuals, the goal is to label anatomically defined regions within bird bodies. This formulation is better suited than instance segmentation for studies that require consistent localisation of intra-organismal structures. Developing such models, however, requires datasets whose labels reflect biological anatomy rather than generic object categories.

A wide range of supervised learning approaches have been developed and applied for part segmentation. Over the past decade, classical architectures such as U-Net (Ronneberger et al., 2015) and DeepLab (Chen et al., 2017; Chen, Papandreou, et al., 2018) have become standard tools for multi-class part segmentation. Specialised network designs have also been proposed specifically for part segmentation, aiming to better capture relationships between object parts and improve fine-grained predictions (Gou et al., 2025; J. He et al., 2023). More recently, open-vocabulary and zero-shot methods, such as VLPart (Sun et al., 2023), have demonstrated the ability to segment directly from text prompts. However, applying these approaches to biological targets can be challenging. Biological part segmentation is domain-specific and often requires anatomically precise definitions that are not well represented in existing computer vision datasets. As a result, supervised models lack standardised benchmarks for systematic evaluation on fine-grained biological tasks, while open-vocabulary and zero-shot methods lack the training signals needed to learn biologically meaningful fine-scale part definitions. These limitations highlight the need for curated, fine-grained datasets to enable both rigorous evaluation and method development for biological part segmentation.

For birds, existing datasets with part-level information remain limited in scope. The CUB-200-2011 dataset provides part annotations as keypoints rather than segmented regions (Wah et al., 2011). PartImageNet (J. He et al., 2022) includes more coarse part segmentations for each object class. For birds, these parts consist of head, body, wing, foot and tail, of which four parts correspond to feathered plumage regions. Therefore, these datasets are insufficient for detailed analyses of plumage structure and colour patterning, and to our knowledge, no existing dataset supports fine-grained, region-level segmentation of avian plumage. Recent advances in vision foundation models (Bommasani et al., 2022)— large-scale models pre-trained on diverse image corpora and designed to transfer across tasks—have opened new opportunities for biological image analysis. Models such as the Segment Anything Model (SAM) (Kirillov et al., 2023) and self-supervised visual transformers such as DINO (Caron et al., 2021), have demonstrated strong transferability across diverse visual domains. In particular, the emergence of larger-scale models such as DINOv3 further suggests the potential for accurate part segmentation with minimal task-specific retraining (Siméoni et al., 2025). Related self-supervised foundation models have already shown strong performance in data-limited medical imaging applications (C. Liu et al., 2025). Recent work has also explored their use for animal part segmentation, for example by applying SAM-based models to segment body parts in fish images (Mehrab et al., 2025). These results suggest that foundation models may enable scalable part segmentation across diverse animal taxa. However, performance on fine-grained anatomical regions in birds and beyond remains largely unexplored. Motivated by these advances, we conduct a systematic evaluation of foundation-model-based approaches alongside existing part segmentation methods for fine-grained segmentation of animal body regions.

In this study, we present a curated dataset of bird images with fine-grained, region-level annotations across nine anatomically defined plumage patches and use it to systematically evaluate a wide range of segmentation architectures, including recent foundation-model encoders. We identify the configuration that yields the highest accuracy for multi-part plumage segmentation and release a ready-to-use model together with a practical guide for applying the workflow to new datasets. We further demonstrate that this approach generalises well to an external part segmentation benchmark including diverse, non-avian taxa and illustrate its application in automated video-based detection, tracking and segmentation. See Figure 1 for an overview of our work.

**Figure 1.**
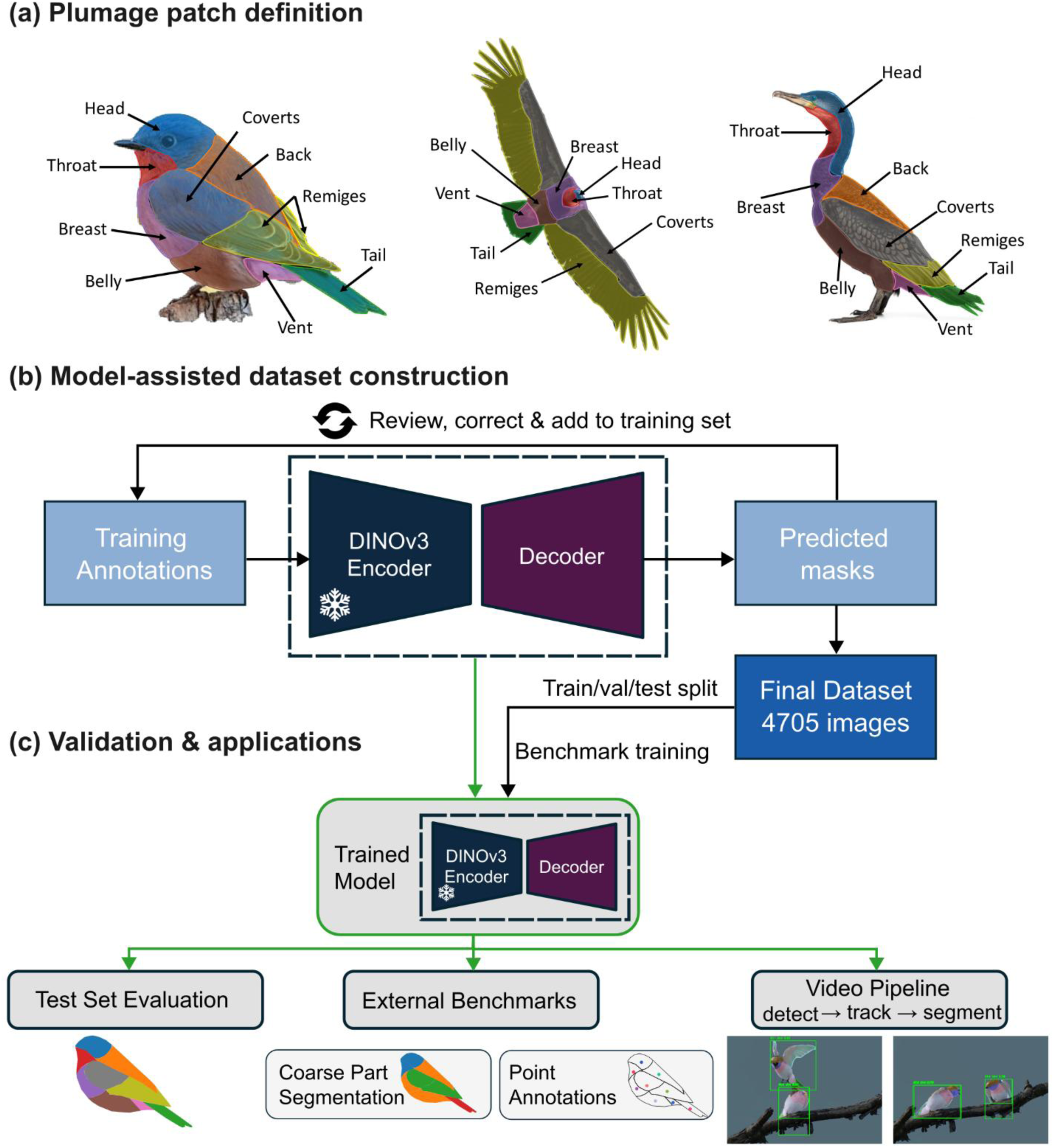
Overview of the plumage patch segmentation workflow. (a) The nine plumage patches defined in this study: head, throat, breast, belly, vent, back, coverts, remiges and tail, shown across three species. (b) Model-assisted dataset construction. Training annotations were used to train a DINOv3-based segmentation model, whose predictions were reviewed, corrected and added back to the training set over successive rounds, yielding a final curated dataset of 4,705 images. (c) Model validation and applications, including held-out test-set evaluation, comparison with external benchmarks providing coarse part segmentations (PartImageNet) or point annotations (CUB-200-2011), and a modular detect–track–segment video pipeline.

## Materials and Methods

### Dataset and Annotations

We selected images from the iRateBirds dataset (Haukka et al., 2023), which compiles bird photographs sourced from the Macaulay Library. Compared with other available avian image datasets, the iRateBirds source dataset offers consistently high-resolution photographs spanning a broad taxonomic range (covering 11,319 species) with diverse backgrounds, making it suitable for developing segmentation models with strong generalisation potential.

Our annotated plumage patch dataset comprised 4,705 images, each representing a unique species, spanning 39 avian orders and 222 families. The dataset was not taxonomically balanced. Passeriformes accounted for the largest proportion of images (60.4%); the number and proportion of images for each order are provided in Table S1. Some images contained multiple individuals; however, all individuals within a given image belonged to the same species. Sex was recorded only from image-level metadata and was not independently validated for each individual; based on these metadata, 408 images were labelled as male, 96 as female and 4,201 as unknown.

Source images were resized so that their longer edge did not exceed 1200 pixels prior to annotation; images with a longer edge already below this threshold were left unresized. Segmentation masks were generated at the corresponding resized resolution. The mean resolution is 1,099 × 793 pixels (most commonly 1,200 × 800 for 1,473 images), with both height and width of at least 400 pixels. Images were selected when most plumage regions were clearly visible, in focus and not substantially obscured by vegetation or other objects.

We defined nine anatomically consistent plumage patches: head, back, tail, throat, breast, belly, vent, coverts and remiges, covering all visible feathered regions (Figure 1a). These definitions follow standard ornithological terminology and were designed to be broadly applicable across taxa, although several anatomical cases required clarification to ensure consistent annotation. The eye region was excluded from the head patch whenever it could be clearly separated; however, when the eye area was small or visually indistinguishable from surrounding plumage, it was included within the head region. In species with extensive bare skin on the head, such as vultures, this region was also annotated as part of the head patch to maintain consistency across taxa. The beak was explicitly excluded from plumage annotations and was not considered part of the head patch. Trousers and undertail coverts were assigned to belly and vent, respectively. Summary statistics of patch coverage and spatial extent are provided in Table S2, and examples illustrating image diversity and representative patch annotations are shown in Figure S1.

To scale annotation efficiently, we used an iterative model-assisted, human-in-the-loop workflow (Figure 1b). An initial DINOv3-based model was trained from 220 manually annotated iNaturalist images, which were used only to bootstrap annotation and were not included in the final dataset. Model predictions were manually inspected and corrected, and the corrected annotations were incorporated into subsequent training iterations. This prediction–correction–retraining process was repeated across five batches of 1,000 images each, until 5,000 images had been reviewed. After quality control, 295 unsuitable images were removed, yielding 4,705 images and 4,781 reviewed bird instances. Of these instances, 60.9% required no correction, 31.6% required minor correction, 6.6% required substantial correction and 0.9% required complete redrawing. Among bird instances requiring manual correction, raw-versus-corrected comparisons showed a mean instance-level mIoU of 0.746. Timing measurements from sampled tasks indicated that the model-assisted workflow was approximately 3.9-fold faster than fully manual annotation (Supplementary Note 1).

To support annotation consistency, reviewers first inspected the raw images independently before viewing the overlaid segmentations, reducing potential confirmation bias from model-generated predictions. Random subsets were cross-checked among annotators to monitor consistency and maintain labelling quality. Images were manually annotated, inspected and corrected by four researchers with expertise in avian anatomy using PhenoLearn (Y. He et al., 2025) and Roboflow (Dwyer et al., 2024). Full details of the annotation workflow, exclusion criteria, correction grades, raw-versus-corrected mIoU, timing estimates and benchmark split composition are provided in Supplementary Note 1, Table S3, Table S4 and Figure S2.

Compared to the CUB-200-2011 and the bird subset of PartImageNet, our dataset prioritises broad taxonomic coverage and detailed plumage characterisation (nine patches). CUB has 11,788 images and covers 200 North American species and provides only landmark points, limiting its utility for morphometric studies requiring area quantification. Similarly, the bird subset of PartImageNet has 2,340 images, offers limited taxonomic coverage (approximately 14 species or superior taxa based on WordNet synsets used in ImageNet, Deng et al., 2009) and coarse granularity (five parts). Image resolutions in CUB-200-2011 and the bird subset of PartImageNet are comparable, with mean resolutions of 468 × 386 pixels (most commonly 500 × 333; Wah et al., 2011) and 492 × 421 pixels (most commonly 500 × 333), respectively. Our dataset thus bridges a critical gap by providing pixel-wise annotations across a phylogenetically diverse range and at a higher resolution. Additionally, given the domain similarity, we also utilise these datasets to evaluate cross-dataset generalisation (see Evaluation section).

### Model Implementation

The best-performing model identified in this study consists of a DINOv3 encoder (ViT-H+/16 distilled variant, approximately 840 million parameters) coupled with a lightweight decoder for dense prediction. We evaluated two decoders inspired by widely adopted segmentation paradigms, U-Net (Ronneberger et al., 2015) and Feature Pyramid Networks (FPN) (Lin et al., 2017), both of which are established and computationally efficient architectures. The first is a Multi-stage Upsampling (MSU) decoder inspired by the U-Net, which gradually increases spatial resolution through a sequence of convolution and upsampling operations. The second is a multi-layer fusion (MLF) decoder inspired by the top-down fusion strategy of FPN, which aggregates feature representations from multiple transformer layers at a fixed spatial resolution. Schematic diagrams of the two decoder designs are provided in Figures S3 and S4. The loss function was defined as a hybrid function combining Dice loss and cross-entropy loss with equal weighting, which is commonly used in biomedical imaging (Isensee et al., 2021). The dataset was split into training, validation and test sets using an 80/10/10 partition (image counts: 3763/470/472). Images and masks were resized with aspect-ratio preservation so that the longest side was 1024 pixels, then padded to 1024 × 1024 pixels for training and evaluation. Images were normalised using ImageNet mean and standard deviation. The model was trained for 50 epochs, with the encoder frozen. Optimisation was performed using the AdamW optimiser with an initial learning rate of 0.001. A linear warm-up was applied over the first 500 optimisation steps, followed by a cosine annealing learning rate schedule.

In addition to the DINOv3-based architectures, we evaluated a range of alternative segmentation models, including U-Net (Ronneberger et al., 2015), DeepLabv3+ (Chen, Zhu, et al., 2018) and SAM-based approaches (Carion et al., 2025; Kirillov et al., 2023), using comparable training settings where applicable. For the SAM-based models, we used only the image encoder paired with the same MSU decoder used for the DINOv3 models. Detailed training configurations and performance summaries for all models are provided in Figure S5. Ablation experiments examining the impact of architectural and training choices for the DINOv3-based models are described in the Ablation Tests section.

### Evaluation

#### Metrics

We primarily evaluated segmentation performance using the mean Intersection over Union (mIoU), which is the standard evaluation metric for part segmentation tasks and is widely used in existing benchmarks (J. He et al., 2022; Sun et al., 2023). Model performance was assessed on the held-out test set of our dataset.

Specifically, we report the macro-averaged mIoU, computed as the unweighted average of IoU scores across all part classes. For each class *c*, the Intersection over Union is defined as

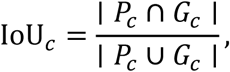

where *P*_*c*_ and *G*_*c*_denote the predicted and ground-truth pixel sets for class *c*, respectively. The mean IoU is then calculated as

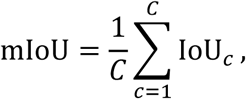

where *C* is the total number of part classes, excluding background.

To evaluate out-of-distribution generalisation, we further tested the best-performing model on two external datasets containing part-level annotations for birds (Figure 1c).

#### Bird Subset of PartImageNet

The first external dataset is the Bird subset of PartImageNet, which contains 2,340 images annotated with five categories: head, body, wing, tail, and foot. As our model specialises in plumage, we established a mapping to aggregate our fine-grained patches into PartImageNet’s parts, as detailed in Table 1. Specifically, predicted masks for constituent patches (e.g., Remiges and Coverts) were merged to form the corresponding coarse region (e.g., Wing). We then computed the mean Intersection over Union (mIoU) between these aggregated predictions and the ground-truth masks.

**Table 1.** Semantic mapping for plumage patches into PartImageNet part categories. Note: The ‘Foot’ category is excluded from evaluation as our model does not predict foot regions.

| **PartImageNet Part** | **Plumage patches** |
| --- | --- |
| **Head** | Head; Throat |
| **Body** | Back; Breast; Belly; Vent |
| **Wing** | Remiges; Coverts |
| **Tail** | Tail |
| **Foot** | No equivalent |

#### CUB-200-2011

We further validated the anatomical correctness of our predictions using the CUB-200-2011 dataset, which contains 11,788 images annotated with 15 anatomical landmarks. To bridge the gap between sparse landmarks and dense segmentation masks, we established a correspondence map linking 10 plumage-relevant landmarks to our patch categories (Table 2). We evaluated spatial consistency by calculating the proportion of landmarks falling within the boundaries of their corresponding predicted plumage regions. For landmarks mapping to multiple patches (e.g., the Nape, which lies on the boundary between Head and Back), a prediction was considered correct if the landmark fell into any valid target region. Non-plumage landmarks (beak, eyes, and legs) were excluded from this analysis. We report the proportion of correctly localised landmarks across all mapped categories.

**Table 2.**
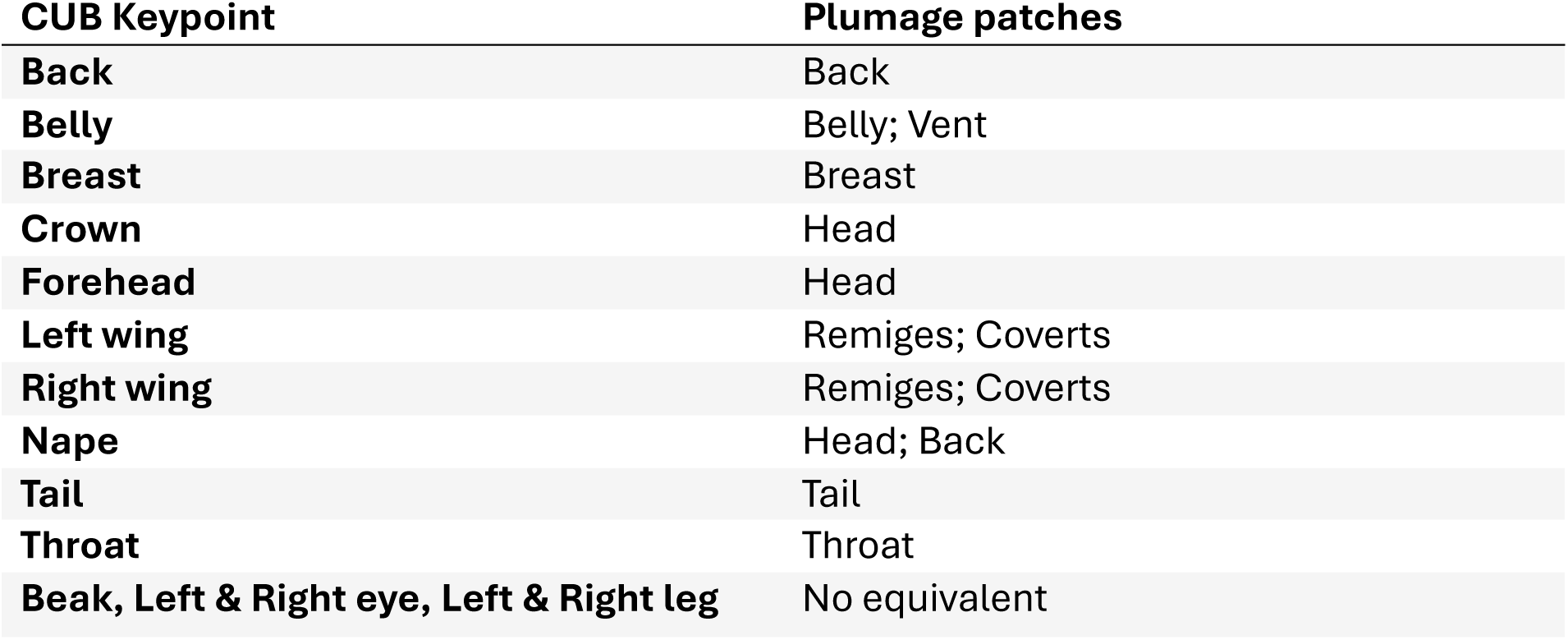
Correspondence between CUB-200-2011 anatomical landmarks and fine-grained plumage patches for spatial consistency evaluation. Note: Beak, eye, and leg landmarks are excluded from evaluation as they correspond to non-plumage regions not predicted by our model.

| CUB Keypoint | Plumage patches |
| --- | --- |
| Back | Back |
| Belly | Belly; Vent |
| Breast | Breast |
| Crown | Head |
| Forehead | Head |
| Left wing | Remiges; Coverts |
| Right wing | Remiges; Coverts |
| Nape | Head; Back |
| Tail | Tail |
| Throat | Throat |
| Beak, Left & Right eye, Left & Right leg | No equivalent |

### Ablation Tests

To assess the contribution of individual design choices in the DINOv3-based model, we conducted a series of ablation experiments in which each variant differed from the baseline in a single component. We first compared Group Normalisation and Batch Normalisation to evaluate optimisation stability under small batch sizes. We then examined the effect of reduced input resolution by training models with smaller spatial dimensions.

We trained models with data augmentation to evaluate the effects of it. Augmentation included bounding-box–based cropping derived from Grounding DINO (S. Liu et al., 2024) bird detections, with all crops manually verified, followed by geometric transformations such as random rotation, flipping and scaling, as well as intensity-based manipulations including sharpening and Gaussian noise.

We further assessed whether the use of class-weighted loss improves performance on smaller plumage patches. Finally, we investigated the role of the DINOv3 encoder by comparing different encoder scales, ViT-B/16 (86 million parameters), ViT-L/16 (300 million parameters) and ViT-H+/16 (840 million parameters). Lastly, we evaluated whether training both the encoder and decoder, rather than freezing the encoder, improves performance on our dataset. All ablation configurations and corresponding results are summarised in Table S5.

### Benchmark on PartImageNet

To assess whether the DINOv3-based architecture performs well beyond our own dataset, we further evaluated it on the supervised part segmentation task of PartImageNet (J. He et al., 2022). PartImageNet spans a broad taxonomic range and is particularly relevant for biological applications, as six of its eleven supercategories represent animals (quadrupeds, bipeds, fish, birds, snakes and reptiles). Performance on this benchmark can provide an indication of the model’s suitability for fine-grained part segmentation across diverse biological forms.

We adopted the official dataset partition, comprising 20,466 training, 1,206 validation, and 2,408 test images, along with the standard part-level annotations. We compared our results with published PartImageNet results, including classical architectures such as DeepLabv3+ (Chen, Zhu, et al., 2018) and more recent approaches, notably the 2025 state-of-the-art method Knowledge-Guided Part Segmentation (KGPS) (Gou et al., 2025), which integrates part-level structural relationships via a large-language-model-derived knowledge graph and a graph convolutional network.

To ensure a fair comparison with KGPS, we matched the training configuration as closely as possible to its published settings. The main comparison used an input resolution of 512 × 512 pixels and no data augmentation. Images and masks were resized with aspect-ratio preservation and padded to the target resolution. The segmentation head was adapted to predict the 41 PartImageNet part classes, while the DINOv3 encoder was kept frozen during training. We evaluated both MSU and MLF decoders. In additional experiments, we explored the effects of higher input resolution (1024 × 1024 pixels) and full end-to-end training in which the DINOv3 encoder was unfrozen. Models were trained using the AdamW optimiser with a learning rate of 0.001 for 30 epochs, and performance was evaluated using mean Intersection over Union (mIoU) on the PartImageNet test set.

### Video Tracking and Segmentation

We further implemented a proof-of-concept pipeline for video-based bird detection, tracking and fine-grained plumage segmentation, using our trained DINOv3 model as the segmentation module. The pipeline supports object detection using either YOLO (for predefined classes, including “bird”) (Redmon et al., 2016) or Grounding DINO (S. Liu et al., 2024), which allows natural-language prompts to specify target objects. Detected objects are linked across frames using SORT (Bewley et al., 2016), after which our model produces multi-class plumage segmentation masks for each detected bird in each frame. The code is designed to be modular, allowing users to replace the detector, tracker or segmentation model for other targets or datasets. This pipeline provides a proof-of-concept route for generating frame-wise plumage segmentations from videos, with potential applications in ecological or behavioural image analysis.

### Software implementation

All models and pipelines developed in this study were implemented in Python. The released codebase provides modular scripts for model training, inference and evaluation, configuration files specifying model architectures and training settings, and an integrated detect–track–segment pipeline for video-based bird analysis. Users can train segmentation models on custom datasets, generate part-level predictions for still images, and apply the framework to videos using detectors, such as YOLO or Grounding DINO. The repository also includes a dataset download and reconstruction script (*download_dataset.py*) that retrieves the corresponding source images from their original Macaulay Library records, subject to the applicable source-image licence and reuse terms.

Model development and training were performed across multiple computing systems. Runtime and memory measurements reported in this study were obtained on a single reference machine to ensure consistency. This system ran Ubuntu 24.04 and was equipped with a single NVIDIA RTX PRO 6000 Blackwell GPU. Experiments were conducted using Python 3.11, PyTorch 2.x and CUDA 12.8.

## Results

### Plumage patch model performance

On the held-out test set (472 images), the best-performing model, based on a frozen DINOv3 encoder with a Multi-stage Upsampling (MSU) decoder, achieved a mean Intersection over Union (mIoU) of 84.01% across all plumage patches, excluding background. Performance varied across anatomical regions (see Table 3), with the highest IoU observed for the head patch (87.5%) and the lowest for the back (75.1%). Other patches with IoU values below 80% included the throat (77.7%) and vent (77.6%).

**Table 3.** Per-class segmentation performance of the best-performing model. IoU scores for each of the nine plumage patch categories evaluated on the held-out test set (n=472). Results are reported for the DINOv3 (ViT-H+/16) encoder with the MSU decoder.

| **Plumage patch** | **IoU (%)** |
| --- | --- |
| **Head** | 87.54 |
| **Back** | 75.06 |
| **Tail** | 85.47 |
| **Throat** | 77.70 |
| **Breast** | 84.37 |
| **Belly** | 84.96 |
| **Vent** | 77.62 |
| **Coverts** | 81.96 |
| **Remiges** | 84.21 |
| **Mean** | **84.01** |

Qualitative inspection of the segmentation results (Figure 2) indicates the model is robust, segmenting multiple individuals within the same image, and remains stable under diverse poses, viewpoints and imaging contexts, including standing, flying and swimming birds. In addition, the model generally avoids segmenting spurious regions arising from reflections, indicating a degree of robustness to visually similar background structures. Most errors arise from intrinsic ambiguity in patch visibility. In particular, patches such as the vent and back are difficult to delineate in certain viewpoints where they are partially or entirely occluded. Boundary uncertainty between neighbouring patches, including the head and throat or remiges and coverts, also contributes to some inconsistencies. These cases typically reflect gradual colour transitions or foreshortening effects rather than clear misclassification. Overall, the qualitative results suggest that the model’s predictions are anatomically coherent and that most errors occur in visually ambiguous situations where precise patch delineation is challenging even for human annotators.

**Figure 2.**
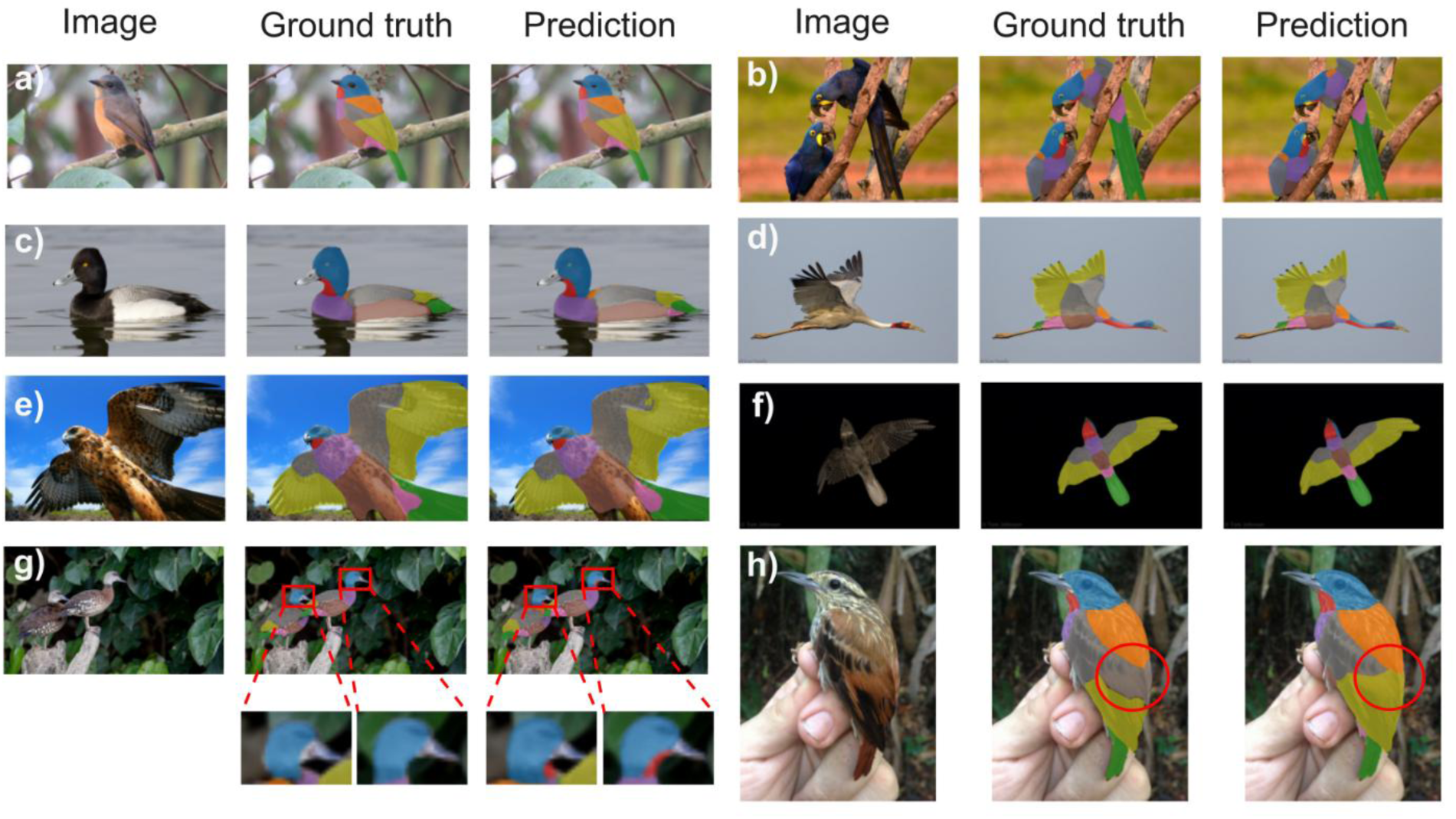
Examples of plumage patch segmentation on the held-out test set. Columns display the input image (left), ground-truth (middle) and prediction (right). (a) A representative example showing clear segmentation of standard anatomical regions. (b) The model successfully segments multiple individuals. (c) Robustness to environmental noise, demonstrating the ability to ignore reflections on the water surface. (d–f) Accurate segmentation of flying birds from (d) side view, (e) ventral view and (f) in low-contrast conditions. (g–h) Examples of prediction ambiguities. (g) Viewpoint ambiguity around the neck-throat region; the model predicts a specific ‘throat’ patch (highlighted in red bounding boxes), whereas the ground truth assigns these pixels to the head and breast. (h) Uncertainty in defining the precise boundary between the back, coverts and remiges (highlighted in red circles).

Results for additional model architectures and training configurations evaluated in this study are summarised in Figure S5. Overall, DINOv3-based models consistently outperformed the other approaches tested. Using an MLF decoder with the DINOv3 encoder resulted in slightly lower performance (mIoU = 82.99%) than the MSU decoder. Classical architectures with ImageNet pretraining, such as DeepLabv3 and U-Net, remained less accurate than DINOv3-based models, suggesting that conventional supervised pretraining was less effective for this fine-grained part segmentation task than self-supervised foundation-model representations. DINOv2-based models achieved competitive performance but did not surpass DINOv3. Segment Anything–based approaches, although also foundation models, performed markedly worse, with mIoU values at least 20 percentage points lower than the best DINOv3-based model. This contrast suggests that foundation-model status alone is insufficient: models trained for whole-object segmentation may not transfer well to biologically defined internal part segmentation. Finally, models trained without weighted loss functions outperformed those using them, indicating that explicit loss re-weighting was not beneficial under our training settings.

#### Bird Subset of PartImageNet

To evaluate cross-dataset generalisation, we applied the model to the bird subset of PartImageNet (2,340 images) using merged predictions corresponding to the four comparable coarse classes (head, body, wing and tail. See Table 1). The resulting mean IoU was 58.7%, with per-class IoUs of 66.6% (head), 48.4% (body), 33.1% (wing) and 47.3% (tail). Detailed class-wise results are provided in Table S6. Qualitative examination suggests that many discrepancies arise from inconsistencies in the PartImageNet annotations, such as some wing regions labelled as body and variable treatment of the beak as part of the head. Consequently, the reported IoU likely reflects both model performance and annotation noise in the benchmark (see Figure 3)

**Figure 3.**
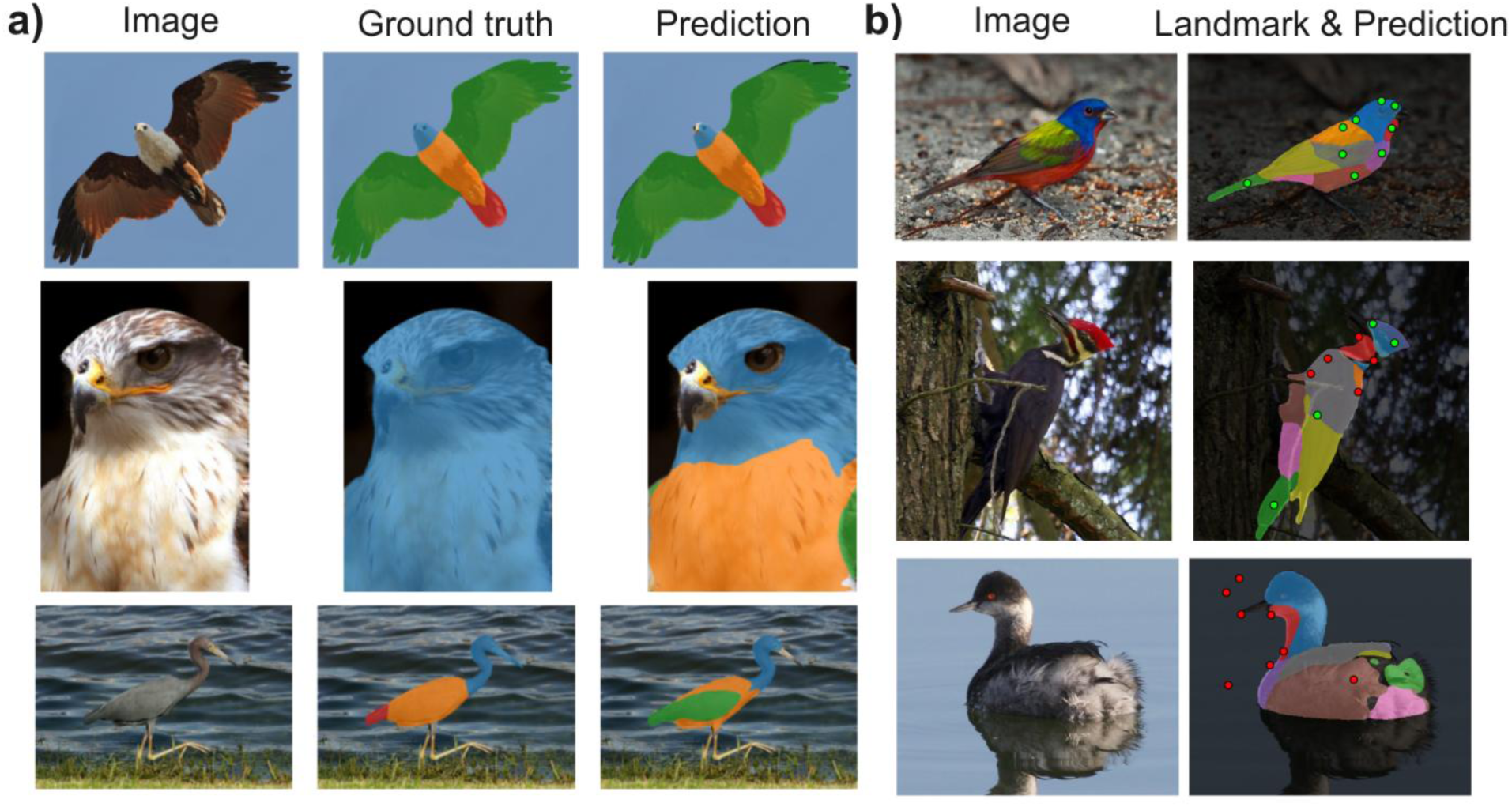
Evaluation of generalisation performance on external datasets. (a) Examples on the PartImageNet bird subset. Columns display the input image, ground-truth and prediction (classes merged to match PartImageNet categories). The top row demonstrates an accurate prediction. However, discrepancies often reflect annotation inconsistencies rather than model failure: the middle row shows the ground truth including the beak, the throat and the breast within the ‘head’ category, while the prediction almost correctly segmented these categories (except the throat); The bottom row highlights ambiguous definitions between wing, tail and body regions. (b) Examples on the CUB-200-2011 dataset. Ground-truth landmarks are overlaid on the model’s predicted segmentation masks. Green dots indicate keypoints correctly falling within the predicted anatomical region; red dots indicate mismatches. The top row shows high spatial agreement. The middle row illustrates ambiguities (e.g. throat/breast, coverts/remiges/back), consistent with results on our holdout test set. The bottom row highlights a case of 0% quantitative accuracy that is actually attributable to a systematic annotation error, where the ground-truth coordinates are shifted along the x-axis relative to the bird. Note that colour schemes differ between panels (a) and (b) due to differences in class definitions. In panel (a), colours correspond to the four merged PartImageNet categories (head, body, wing and tail), whereas in panel (b), colours follow the nine plumage patch classes defined in this study. In both panels, colours are assigned according to class index using the same colour palette (Tab10).

#### CUB-200-2011

We further evaluated spatial consistency using the CUB-200-2011 dataset, which provides keypoint annotations rather than segmentation masks. Across 11,788 images and 100,718 valid keypoints, 93,520 points fell within the corresponding predicted plumage patches, yielding an overall localisation accuracy of 92.9%. Except for the back keypoint (79.2%), all other mapped keypoints achieved localisation accuracies above 85%. Detailed per-keypoint accuracy is reported in Table S7. Inspection of low-scoring cases revealed that some apparent failures were attributable to errors in the CUB annotations themselves. For example, several images with 0% keypoint accuracy exhibited systematic shifts of keypoint coordinates away from the bird, while the predicted plumage segmentations remained anatomically plausible (Figure 3).

### Ablation Tests

Ablation experiments revealed distinct effects of data-related designs and architectural configurations on segmentation performance (Table S5). Reducing the input resolution led to a consistent decrease in accuracy, with a drop of more than 10 percentage points at 256 × 256 pixels, indicating that fine-grained plumage segmentation strongly benefits from higher spatial resolution. The inclusion of geometric and intensity-based data augmentation did not improve test-set performance and led to a small decrease in mIoU (approximately 1– 2 percentage points), suggesting limited benefit of such augmentations for this task.

The model remained relatively robust under reduced training data availability. When trained on approximately 10% of the original training set, performance decreased by around 5 percentage points. Even under more extreme data reduction, the model achieved an mIoU of 74.63% when trained on 37 images and 66.76% when trained on only 6 images. These results demonstrate the model’s capability to learn effective segmentation features from extremely limited supervision.

Although batch normalisation typically requires large batch sizes to achieve stable optimisation, we also observed this effect when training DeepLabv3 models from scratch. Replacing group normalisation with batch normalisation in the DINOv3-based model resulted in only minor differences in performance (less than 1 percentage point) under our training settings.

Architectural ablations indicated that reducing the DINOv3 encoder size led to a modest but consistent decrease in accuracy, suggesting that larger encoders increase performance. In contrast, jointly training the entire model by unfreezing the DINOv3 encoder resulted in a substantial drop in performance (84.01 vs 66.11; approximately 18 percentage points) compared with keeping the encoder frozen. Full fine-tuning also had a higher memory cost, with GPU memory usage increasing from approximately 6.6 GB when training only the decoder to 39 GB when training the entire model.

### Performance on PartImageNet

On the PartImageNet benchmark, our DINOv3-based model performed competitively with previously published methods (Table 4). Using a frozen DINOv3 encoder with an MLF decoder and a 512 × 512 input resolution, the model reached an mIoU of 73.17%, comparable to the 72.69% reported for KGPS at matched resolution (512×512).

**Table 4.** Performance on the PartImageNet benchmark test set (N=2408). Models are compared using mean Intersection over Union (mIoU) on the official PartImageNet test split. For fair comparison with other models, our runs were trained without data augmentation. “Frozen” indicates that the DINOv3 encoder was not fine-tuned.

| Model | Input resolution | Encoder training | mIoU (%) |
| --- | --- | --- | --- |
| **DeepLabv3+** | $512 \times 512$ | – | 60.57 |
| **Compositor** | $512 \times 512$ | – | 64.64 |
| **CAT-Seg** | $512 \times 512$ | – | 67.73 |
| **KGPS (2025)** | $512 \times 512$ | – | 72.69 |
| **DINOv3 + MSU** | 512 × 512 | Frozen | 72.86 |
| **DINOv3 + MLF** | 1024 × 1024 | Frozen | 76.11 |
| **DINOv3 + MLF** | 1024 × 1024 | Full training | 71.04 |
| **DINOv3 + MLF** | **512 × 512** | **Frozen** | **73.17** |

Increasing the input resolution to 1024 × 1024 pixels further improved performance, yielding an mIoU of 76.11%. In contrast, full end-to-end training in which the DINOv3 encoder was unfrozen resulted in lower performance (71.04%), suggesting that fine-tuning such a large foundation model on PartImageNet may require more careful optimisation or larger training data.

The model with MSU decoder, which performed best on our plumage patch dataset, achieved an mIoU of 72.86% on PartImageNet, indicating that the approach generalises well across datasets. Overall, these results demonstrate that a simple decoder built on top of a frozen DINOv3 encoder is competitive on a challenging multi-class part segmentation benchmark.

In addition to quantitative evaluation, PartImageNet allows a qualitative assessment of model performance across a range of non-avian animal taxa, including quadrupeds, fish, snakes and reptiles. Figure 4 shows representative examples of part segmentations produced by our DINOv3-based model on these categories. Although PartImageNet provides only coarse part annotations that do not capture fine-grained anatomical subregions, the model recovers coherent and anatomically plausible part structures across diverse organisms. These results provide a preliminary indication that the proposed approach is not limited to avian morphology and can be transferred to other taxa, given appropriate part definitions and annotation schemes.

**Figure 4.**
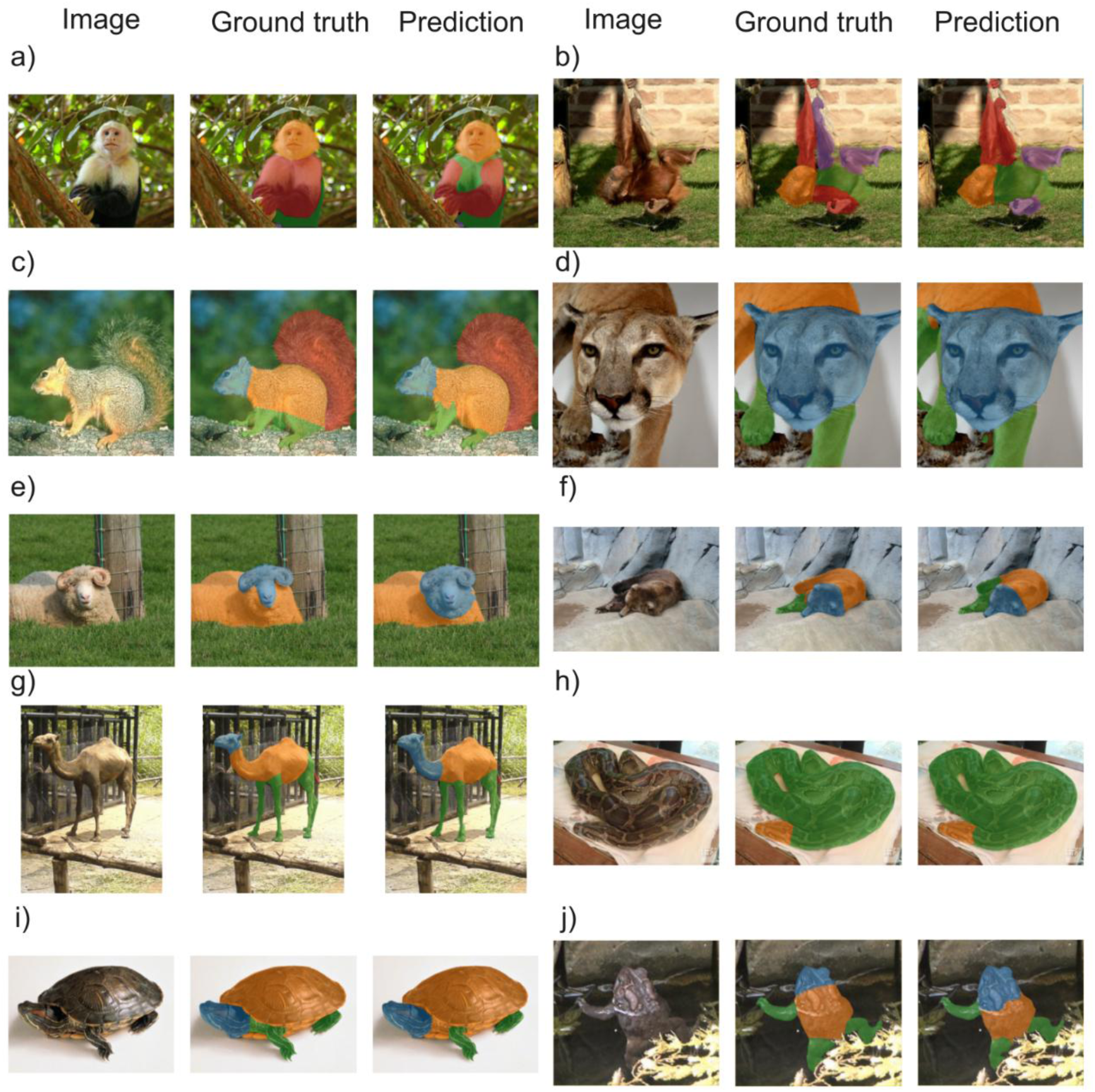
Generalisation of the segmentation model to diverse non-avian taxa on the PartImageNet benchmark. 10 representative examples illustrate the model’s ability to adapt to distinct morphological structures beyond birds. Each panel displays a triplet containing the original image (left), ground-truth annotation (centre) and model prediction (right). The selection covers a broad taxonomic range, including (a-g) mammals (primates, rodents, carnivores and ungulates), (h, i) reptiles (snakes and turtles) and (j) amphibians.

### Video Tracking and Segmentation

We applied our detection–tracking–segmentation pipeline to a small set of bird videos to demonstrate its applicability in dynamic contexts. Because ground-truth multi-part video annotations were unavailable, we did not perform quantitative benchmarking on video data. Instead, qualitative inspection of example outputs indicated visually coherent identity tracking and multi-part plumage segmentation across frames (see Figure S6).

To provide an indication of computational performance, we report average per-frame runtimes for a representative 1920 × 1080 video containing two birds. Mean detection time using Grounding DINO was 8 × 10⁻³ s per frame, tracking with SORT required 2.5 × 10⁻⁵ s per frame, and multi-part segmentation required 0.69 s per frame when using the DINOv3 ViT-H+/16 encoder (the best-performing variant). Using the smaller DINOv3 ViT-B/16 encoder, which has approximately one-tenth the number of parameters, reduced segmentation time to 0.27 s per frame. These measurements are intended to illustrate relative computational costs across model variants rather than to provide a benchmark.

## Discussion

Fine-grained characterisation of avian plumage patches is central to many questions in ecology and evolutionary biology, including sexual ornamentation, sexual dichromatism and ecological variation in colouration across body regions (Cooney et al., 2019, 2022; Dale et al., 2015; López-Idiáquez et al., 2022; Nolazco et al., 2023; Shultz & Burns, 2017). Progress in these areas has been constrained partly by the absence of image resources with annotations grounded in biologically meaningful patch definitions. Existing datasets designed for computer vision tasks offer limited utility in this context: the bird subset of PartImageNet (J. He et al., 2022) uses five coarse part categories (head, body, wing, tail and foot), and as illustrated in Figure 3a, the assignment of plumage regions to these categories varies across species in ways that do not reflect consistent anatomical criteria. CUB-200-2011 (Wah et al., 2011), while biologically motivated, records part locations as keypoints that cannot support region-level analyses such as patch area quantification or texture measurement. Our dataset addresses these limitations directly. The nine patch categories follow standard ornithological terminology, providing semantically consistent, pixel-level boundary annotations that are immediately interpretable in a biological context. The dataset offers broader phylogenetic coverage than either PartImageNet or CUB, and images were acquired at a higher average resolution than either dataset. Together, these properties position the dataset as a resource that bridges the gap between the scale of computer vision benchmarks and the biological precision required for ecological and evolutionary research.

One immediate potential application is research on visual perception and aesthetics. The iRateBirds source corpus (Haukka et al., 2023) was designed to elicit human aesthetic responses to bird photographs, and our patch-level annotations enable decomposition of these holistic responses into patch-level contributions, such as identifying which plumage regions most strongly predict aesthetic ratings or how patch-level colour and pattern variation shape perceptual judgements. Because both the patch features and the perceptual ratings derive from the same set of photographs, such analyses remain consistent under variable photographic conditions and do not require colour calibration, contributing to a growing literature linking human aesthetic perception to broader questions in evolutionary biology and signalling ecology (Renoult & Mendelson, 2019).

The dataset also addresses a resource gap for computer vision research on biologically grounded animal recognition. Existing avian benchmarks provide relatively coarse part categories, as in PartImageNet (head, body, wing, tail and foot). Our nine-category ornithological annotation may therefore serve as a benchmark for developing and evaluating segmentation, recognition, and open-vocabulary approaches that aim to align visual representations with biologically meaningful anatomical concepts (Sun et al., 2023). The annotations may also support the development and evaluation of interpretable fine-grained classification methods (Kumar & Kondaveeti, 2024), where part-level information can be used to inspect whether classifiers attend to biologically meaningful regions.

Our results demonstrate that accurate fine-grained part segmentation can be achieved across a broad taxonomic sample of birds and may provide a useful pretraining source for more controlled phenotyping datasets. Direct measurement of colour requires colour-calibrated conditions and controlled lighting (e.g., Y. He et al., 2022), while geometric morphometric analyses require consistent image acquisition and viewing geometry to ensure that landmark or outline coordinates are comparable across specimens (e.g., Zelditch et al., 2012). Both requirements are typically met through museum specimen photography or dedicated field protocols. Although our dataset lacks the calibration needed for direct quantitative phenotyping, it can provide a pretraining source for models applied to controlled-photography datasets. Because such datasets often contain less background heterogeneity and more standardised imaging conditions than naturalistic imagery, models pretrained on our data may provide a strong starting point, although target-domain fine-tuning will likely be required.

Beyond its analytical use, accurate part segmentation also has direct value for dataset construction itself. Region-level part segmentation is costly to annotate manually: unlike image classification or object detection, which require only image-level labels or bounding boxes, and unlike whole-object segmentation, which delineates a single outline, part segmentation requires multiple defined regions within each individual. In practical human-in-the-loop workflows, stronger models can generate more accurate initial masks and thereby reduce the manual correction required to build or extend annotated datasets. In our workflow, this reduced annotation effort: the model-assisted procedure was approximately 3.9-fold faster than annotating from scratch (Supplementary Note 1), allowing annotators to refine predicted masks rather than draw each region manually.

Our benchmarking further shows that accurate segmentation can be achieved with limited computing and data: fine-grained segmentation performance improves substantially when using DINOv3 compared with classical convolutional architectures. One likely explanation is that DINOv3 provides intermediate representations that jointly capture strong semantic information and high-resolution structural detail (Siméoni et al., 2025), both of which are critical for anatomically meaningful part segmentation. This does not imply that any foundation model will suffice. SAM-based encoders performed poorly on our task, likely because SAM was originally designed for prompt-based segmentation of whole objects with relatively well-defined boundaries rather than for delineating fine-grained regions. However, this comparison should be interpreted cautiously. We evaluated SAM through its image encoder rather than its full prompt-based architecture; a complete assessment of prompt-based SAM variants adapted for dense part prediction is a worthwhile direction for future work.

As noted above, our dataset annotations were produced through a DINOv3-assisted procedure. Because DINOv3-based models also performed best in our benchmark, one possible concern is that the evaluation is biased toward predictions resembling the assisting model. Several considerations mitigate this concern. The assisting and evaluated models were trained separately, and although 60.9% of instances required no correction, these were predominantly visually unambiguous individuals for which predictions were already correct rather than cases of uncritical acceptance; annotators inspected each raw image before viewing predictions, and the remaining 39.1% received manual correction (raw-versus-corrected mIoU of 0.746). In addition, our DINOv3-based models also achieved strong performance on the full PartImageNet benchmark (Table 4), whose annotations are entirely independent of our pipeline and therefore cannot be affected by any such circularity. Consistent with this, models trained on our dataset generalised well to the independently annotated PartImageNet bird subset and CUB-200-2011. Together, these results indicate that the advantage of DINOv3-based representations is unlikely to be explained solely by the annotation process.

A practical implication of our benchmarking is that freezing the foundation model encoder and training only a lightweight decoder outperformed end-to-end fine-tuning, while requiring less than 20% of the GPU memory (6.6 GB vs 39 GB). A likely explanation is that our dataset (∼4,000 images) is insufficient to stably fine-tune a model with approximately 840 million parameters, and effective optimisation of such large models typically requires extensive hyperparameter exploration. We note, however, that this finding may reflect our specific fine-tuning configuration; systematic exploration of learning-rate schedules and gradual layer-wise unfreezing could yield different conclusions and remains an important direction for future work. Several further findings reinforce the accessibility of this approach: ablation experiments showed that commonly used techniques such as data augmentation and class-weighted loss did not improve performance, suggesting that DINOv3 features can distinguish minor classes without augmentation or reweighting, and the model retained reasonable performance when trained on severely reduced subsets of the data. These results suggest that a frozen encoder with a lightweight decoder provides a robust and efficient starting point for fine-grained biological part segmentation, lowering the computational and technical barriers to adopting AI-based image analysis in ecological and evolutionary research (Bianchini et al., 2025).

Taken together, our work demonstrates that fine-grained biological part segmentation can be achieved with relatively simple training procedures, a single GPU and moderate annotated data, and that model-assisted annotation can help scale the construction of such datasets. Beyond birds, the same frozen-encoder approach achieved good performance on diverse non-avian taxa in PartImageNet, indicating that the method can extend to broader animal part-segmentation tasks given appropriate training annotations (Figure 4). By releasing the dataset, code and trained model, we provide a practical resource for ecology and evolutionary biology groups seeking to incorporate part segmentation into avian image-analysis workflows, or to extend it to other animal taxa. More broadly, our results point to robust foundation-model encoders, combined with open benchmark datasets and modular analysis pipelines, as a practical route for scaling biologically meaningful image segmentation.

## Data and code availability

The PlumageParts annotation masks, metadata, train/validation/test split information, prediction masks and trained model checkpoints are available from Zenodo: https://doi.org/10.5281/zenodo.20551408. Original source images are not redistributed in this release because they remain subject to the applicable Macaulay Library source-image licence and reuse terms. To support reproducibility, the released metadata includes the corresponding Macaulay Library record information, and the GitHub repository provides a *download_dataset.py* script to help users retrieve the source images and reconstruct the dataset locally, subject to those terms. The source code, documentation and dataset reconstruction instructions are available at https://github.com/EchanHe/PlumageParts.

## Acknowledgements

We thank the Macaulay Library at the Cornell Lab of Ornithology for confirming permission to use source photographs for the creation of segmentation annotations and for their display in this manuscript. We thank C. Niu for assistance with the project and J. Han for valuable comments on the project. We are also grateful to our colleagues and friends at the University of Sheffield for their support during the course of this work.

## Author contributions

Y.H.: Conceptualisation, Methodology, Software, Data curation, Analysis, Visualisation, Writing — original draft.

E.I.: Conceptualisation, Methodology, Software, Analysis, Visualisation, Writing — review & editing.

K. H: Methodology, Data curation, Visualisation, Writing — review & editing. G.T.: Supervision, Writing — review & editing.

S.M.: Supervision, Methodology, Writing — review & editing.

J.P.R.: Supervision, Methodology, Data curation, Writing — review & editing.

C.C.: Supervision, Methodology, Analysis, Writing — review C editing.

## Declaration of generative AI

During the preparation of this work, the authors used generative AI to assist with grammar checking. After using this tool, the authors reviewed and edited the content as needed and took full responsibility for the content of the article.

