## Supplementary information for "PlumageParts: A fine-grained avian plumage segmentation dataset and benchmark for ecological image analysis"

### Supplementary Note 1. Model-assisted annotation workflow, quality control and correction effort

#### Iterative model-assisted annotation workflow

To segment plumage patches efficiently, we used an iterative model-assisted annotation workflow. We first trained an initial segmentation model using 220 manually annotated bird images selected from iNaturalist. These images were segmented by the authors and were used only to bootstrap the annotation-assistance workflow; they were not included in the final curated dataset or in the benchmark splits. Following a preliminary qualitative comparison of candidate segmentation models, we selected a DINOv3 encoder with a lightweight decoder as the annotation-assistance model.

The initial model was used to generate predictions for the first batch of 1,000 target bird images. The predicted masks were manually inspected and corrected, and the resulting annotations were then added to the training set to train the next model iteration. This process was repeated in batches of 1,000 images until predictions had been generated and reviewed for 5,000 target images.

#### Manual quality control

Following a manual quality check, 295 images were excluded, resulting in a final curated dataset of 4,705 images. Images were removed when they were unsuitable for reliable fine-grained plumage annotation, primarily due to insufficient target size or visibility, image blur or poor resolution, poor illumination or low contrast, substantial occlusion, incomplete anatomical coverage, invalid or non-photographic targets, or ambiguous plumage-region boundaries.

Following segmentation prediction, quality was assessed at the individual-bird level rather than at the image level. Although the retained images contained more birds in total, only individuals for which plumage segmentation was feasible were included in the annotation review, yielding 4,781 bird instances. Among the 4,705 images, 4,643 contained one reviewed bird instance, 51 contained two, eight contained three, and three contained four.

Each bird instance was assigned a four-level correction grade: grade-1 indicated an acceptable segmentation requiring no correction; grade-2 indicated minor issues, such as small boundary refinements; grade-3 indicated substantial errors requiring correction of multiple regions or partial redrawing; and grade-4 indicated failed segmentations requiring complete redrawing. Overall, 2,912 bird instances were assigned grade-1, 1,511 grade-2, 315 grade-3 and 43 grade-4, corresponding to 60.9%, 31.6%, 6.6% and 0.9% of reviewed instances, respectively.

**Raw prediction versus corrected annotation agreement**

To quantify the extent of manual correction introduced during model-assisted annotation, we compared the raw model outputs with the final corrected annotations using image-level mean intersection-over-union (mIoU). Across the 4,705 retained images, the mean raw-versus-corrected mIoU was 0.903. The first batch, generated using the bootstrap model trained from the 220 external images, had the lowest mean mIoU of 0.836. After corrected target-domain annotations were incorporated into training, agreement increased and remained broadly stable across subsequent batches, with mean mIoU values of 0.922, 0.921, 0.924 and 0.912 for the subsequent four batches, respectively (See Table S3). In addition, the first annotation batch contained more bird instances requiring manual correction than subsequent batches. In Batch 1, 599 grade-2 to grade-4 instances were recorded, whereas later batches contained 292–352 such instances.

We also summarised raw-versus-corrected agreement for bird instances that required manual correction. Because correction grades were assigned at the bird-instance level, this analysis used instance-level mIoU rather than image-level mIoU. Among grade-2, grade-3 and grade-4 instances, the overall mean mIoU was 0.746. Agreement decreased with increasing correction grade: grade-2 instances had a mean mIoU of 0.780, grade-3 instances had a mean mIoU of 0.645 and grade-4 instances had a mean mIoU of 0.269. The distributions of instance-level mIoU across batches and correction grades are shown in Figure S2.

**Annotation-time estimate**

To estimate annotation efficiency, we measured the time required for annotation and correction tasks using 10 sampled images for each correction-grade category. Because correction grades were assigned at the bird-instance level, we selected single-instance images containing bird instances of the corresponding grade for this timing analysis. Visual inspection of predictions requiring no correction took an average of 5 s per instance. Grade-2 inspection and correction took 32 s on average, grade-3 inspection and correction took 1 min 20 s, and grade-4 re-segmentation took 2 min 34 s. Full manual annotation from scratch, estimated using images containing grade-1 to grade-3 instances, took 1 min 17 s per instance on average.

For grade-1 to grade-3 instances, which accounted for 4,738 of the 4,781 reviewed bird instances, combining these timing estimates with the observed correction-grade distribution gave an expected model-assisted annotation time of 18.6 s per instance, compared with 77 s for full manual annotation from scratch. This corresponds to an approximately 4.1-fold increase in annotation speed. Grade-4 instances were treated separately because they represented failed predictions requiring re-segmentation. For these cases, the model-assisted workflow required visual inspection followed by re-segmentation, giving an estimated time of 159 s per instance, compared with 154 s for manual re-segmentation alone. Across all 4,781 reviewed bird instances, including grade-4 cases, the expected model-assisted annotation time was 19.9 s per instance, compared with 77 s under the corresponding full manual workflow, corresponding to an approximately 3.9-fold increase in annotation speed. Because these measurements were based on 10 sampled images per category rather than a controlled user study, they should be interpreted as an approximate estimate of annotation efficiency.

**Benchmark split**

For benchmarking, the final dataset was randomly divided into training, validation and test splits. The training split contained 3,763 images and 3,823 bird instances, of which 61.0% were grade-1, 31.5% grade-2, 6.6% grade-3 and 0.8% grade-4. The validation split contained 470 images and 476 instances, with 61.6%, 31.7%, 5.7% and 1.1% assigned to grades 1–4, respectively. The test split contained 472 images and 482 instances, with corresponding proportions of 59.3%, 32.2%, 7.3% and 1.2% (Table S4).

**Table S1. Taxonomic composition of the annotated plumage patch dataset.** The table summarises the number of images per avian order in the dataset (4,705 images).

| Order | Count | Ratio |
|---|---|---|
| **Passeriformes** | 2840 | 60.4 |
| **Caprimulgiformes** | 249 | 5.3 |
| **Charadriiformes** | 180 | 3.8 |
| **Piciformes** | 179 | 3.8 |
| **Columbiformes** | 153 | 3.3 |
| **Psittaciformes** | 147 | 3.1 |
| **Accipitriformes** | 114 | 2.4 |
| **Galliformes** | 112 | 2.4 |
| **Strigiformes** | 96 | 2 |
| **Coraciiformes** | 82 | 1.7 |
| **Anseriformes** | 78 | 1.7 |
| **Cuculiformes** | 63 | 1.3 |
| **Gruiformes** | 61 | 1.3 |
| **Procellariiformes** | 60 | 1.3 |
| **Pelecaniformes** | 54 | 1.1 |
| **Bucerotiformes** | 33 | 0.7 |

| | | |
|---|---|---|
| **Falconiformes** | 33 | 0.7 |
| **Trogoniformes** | 28 | 0.6 |
| **Suliformes** | 27 | 0.6 |
| **Galbuliformes** | 27 | 0.6 |
| **Tinamiformes** | 15 | 0.3 |
| **Otidiformes** | 13 | 0.3 |
| **Podicipediformes** | 13 | 0.3 |
| **Musophagiformes** | 10 | 0.2 |
| **Pterocliformes** | 7 | 0.1 |
| **Ciconiiformes** | 6 | 0.1 |
| **Sphenisciformes** | 5 | 0.1 |
| **Phoenicopteriformes** | 4 | 0.1 |
| **Coliiformes** | 3 | 0.1 |
| **Casuariiformes** | 2 | 0 |
| **Apterygiformes** | 2 | 0 |
| **Struthioniformes** | 2 | 0 |
| **Cariamiformes** | 1 | 0 |
| **Cathartiformes** | 1 | 0 |
| **Gaviiformes** | 1 | 0 |
| **Leptosomiformes** | 1 | 0 |
| **Mesitornithiformes** | 1 | 0 |
| **Rheiformes** | 1 | 0 |
| **Eurypygiformes** | 1 | 0 |

**Table S2. Annotation coverage and spatial statistics of plumage patches.** Summary of label completeness and structural variation across the dataset (4,705 images). The table shows the frequency of occurrence and spatial extent (median pixel area and interquartile range) for each of the nine defined plumage regions. High coverage was observed across all categories (ranging from 77.0% for the vent to 99.9% for the head). The substantial variation in pixel area reflects differences in bird species, pose, camera viewpoint, and image scale across the collected photographs.

| Patch | Images (n) | Presence (%) | Median pixels | Interquartile Range pixels |
|---|---|---|---|---|
| **Head** | 4,699 | 99.9 | 9,796 | 5,808–15,948 |
| **Back** | 3,865 | 82.1 | 4,615 | 1,402–11,029 |
| **Tail** | 4,317 | 91.8 | 6,155 | 2,971–11,530 |
| **Throat** | 4,634 | 98.5 | 2,392 | 1,302–4,180 |
| **Breast** | 4,550 | 96.7 | 7,916 | 3,913–14,361 |
| **Belly** | 4,447 | 94.5 | 11,096 | 5,531–19,647 |

| | | | | |
|---|---|---|---|---|
| **Vent** | 3,623 | 77.0 | 3,563 | 1,614–7,052 |
| **Coverts** | 4,623 | 98.3 | 10,022 | 5,134–18,808 |
| **Remiges** | 4,396 | 93.4 | 8,318 | 3,758–15,812 |

**Table S3. Raw-versus-corrected image-level mIoU by annotation batch.** Mean mIoU was calculated between raw model outputs and final corrected annotations for each retained image. The number of grade-2 to grade-4 instances indicates the number of bird instances requiring manual correction in each annotation batch.

| Batch | Images | Grade 2–4 instances | Mean mIoU |
|---|---|---|---|
| **1** | 949 | 599 | 0.836 |
| **2** | 943 | 352 | 0.922 |
| **3** | 939 | 292 | 0.921 |
| **4** | 937 | 311 | 0.924 |
| **5** | 937 | 315 | 0.912 |

**Table S4. Dataset split and correction-grade distribution.** Number of images, bird instances and correction grades in the training, validation and test splits. Percentages indicate the proportion of bird instances assigned to each grade within each split.

| Split | Images | Bird instances | Grade-1 | Grade-2 | Grade-3 | Grade-4 |
|---|---|---|---|---|---|---|
| **Train** | 3,763 | 3,823 | 2,333 (61.0%) | 1,205 (31.5%) | 253 (6.6%) | 32 (0.8%) |
| **Validation** | 470 | 476 | 293 (61.6%) | 151 (31.7%) | 27 (5.7%) | 5 (1.1%) |
| **Test** | 472 | 482 | 286 (59.3%) | 155 (32.2%) | 35 (7.3%) | 6 (1.2%) |
| **Overall** | 4,705 | 4,781 | 2,912 (60.9%) | 1,511 (31.6%) | 315 (6.6%) | 43 (0.9%) |

**Table S5. Ablation tests for the DINOv3-based plumage segmentation model.** Unless otherwise stated, each ablation modifies a single component of the baseline model, which consists of a frozen DINOv3 ViT-H+/16 encoder with a Multi-stage Upsampling (MSU) decoder trained at 1024 × 1024 resolution using group normalisation and a hybrid Dice and cross-entropy loss (batch size = 2). All models were trained for 50 epochs using the AdamW optimiser with an initial learning rate of 0.001, a linear warm-up over the first 500 optimisation steps, and a cosine annealing learning rate schedule. Performance is reported as mean Intersection over Union (mIoU) on the held-out test set.

| Test category | Configuration | mIoU (%) |
|---|---|---|
| **Baseline** | ViT-H+/16, 1024×1024,<br>Full training set (3763 images)<br>Group normalisation,<br>No augmentation, frozen encoder | 84.01 |
| **Encoder size** | ViT-B/16 | 77.9 |
| | ViT-L/16 | 81.49 |
| **Input resolution** | 256 × 256 | 71.16 |
| | 512 × 512 | 80.99 |
| **Training set size** | 0.2% (6 images) | 66.76 |
| | 1% | 74.63 |
| | 10% | 79.53 |
| | 20% | 81.03 |
| | 50% | 82.24 |
| **Augmentation** | With augmentation | 82.46 |
| **Normalisation** | Batch normalisation | 83.4 |
| **Encoder training** | Full fine-tuning | 66.11 |

**Table S6. Class-wise segmentation IOU on the bird subset of PartImageNet**.

| PartImageNet part | IOU |
|---|---|
| **Background** | 98.08 |
| **Head** | 66.65 |
| **Body** | 48.43 |
| **Wing** | 33.05 |
| **Tail** | 47.3 |

**Table S7. Per-point accuracy on CUB-200-2011 dataset.**

| CUB Keypoint | Accuracy |
|---|---|
| **Back** | 79.19 |
| **Belly** | 85.59 |
| **Breast** | 94.82 |

| | |
|---|---|
| **Crown** | 98.58 |
| **Forehead** | 97.64 |
| **Left wing** | 95.48 |
| **Right wing** | 95.36 |
| **Nape** | 98.15 |
| **Tail** | 89.75 |
| **Throat** | 92.48 |

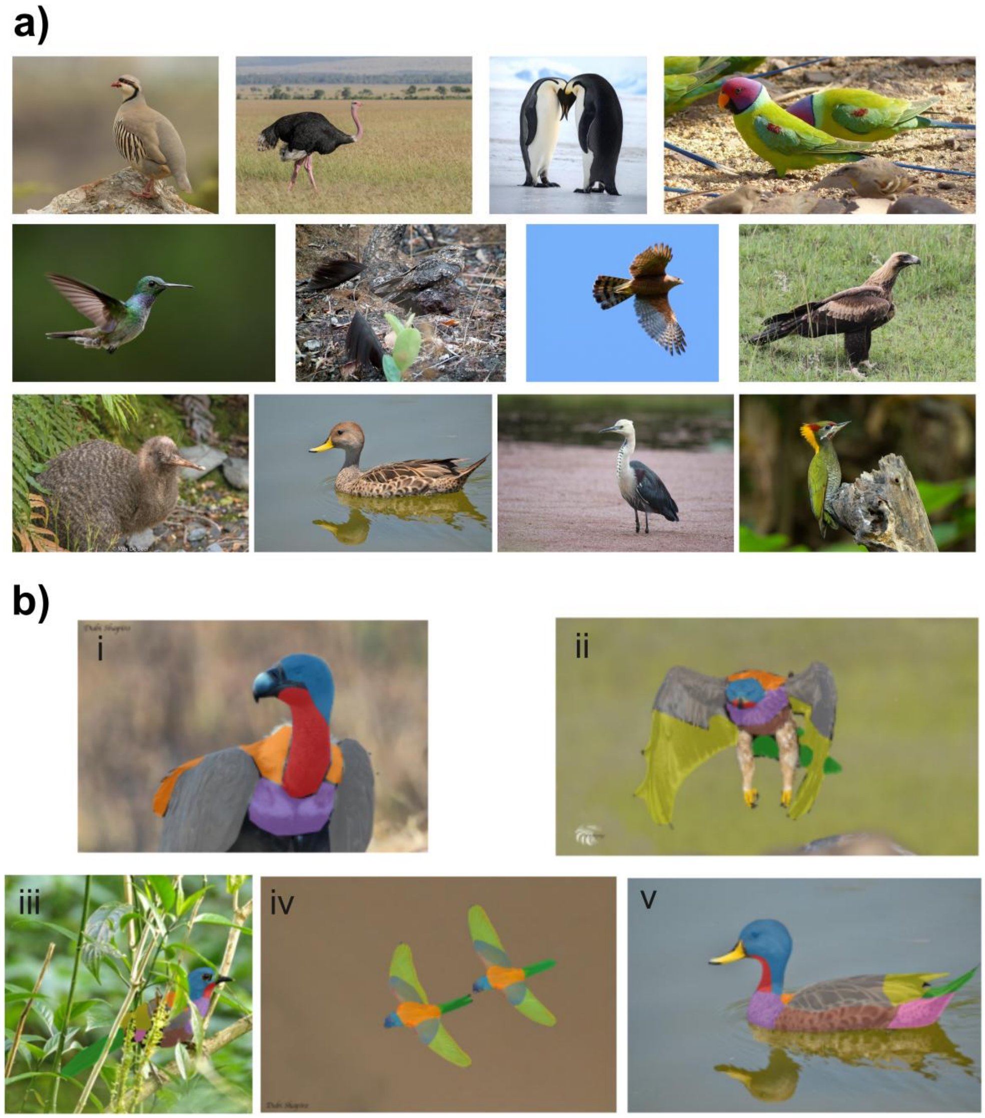


**Figure S1. Dataset diversity and annotation protocols.** (a) Representative examples illustrating the wide variance in species, plumage patterns, poses, and imaging environments (e.g., sky, water, vegetation). (b) Examples of expert labelling decisions in challenging scenarios: (i) Bare skin: For species with bare heads (e.g., vultures), the skin is annotated as the 'head' region to maintain topological consistency. (ii) Feathered legs and claws are excluded from the annotation. (iii) Occlusion handling: In cases of partial occlusion, segmentation masks label visible plumage only, excluding foreground objects (e.g., leaves). (iv) Multi-instance labelling: All visible individuals within an image are annotated. (v) Reflections on water or other surfaces are excluded from the segmentation masks.

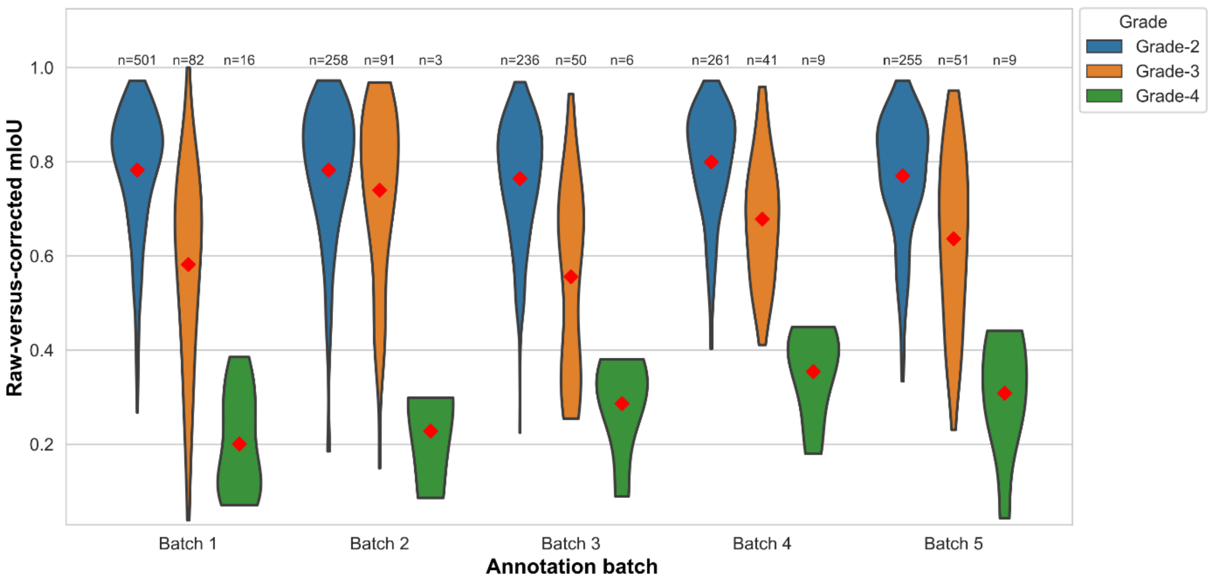


**Figure S2. Raw-versus-corrected agreement by annotation batch and correction grade.** Violin plots show instance-level mIoU between raw model predictions and final corrected annotations for grade-2, grade-3 and grade-4 bird instances. Results are grouped by annotation batch. Red diamonds indicate mean mIoU, and labels above each violin indicate the number of instances. Higher correction grades correspond to larger manual modifications.

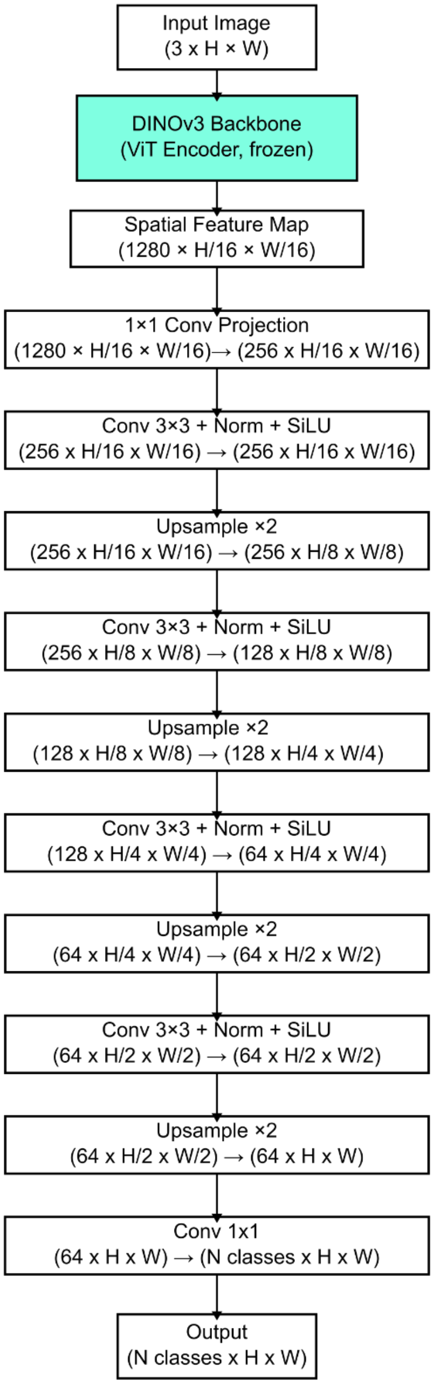


**Figure S3. Multi-stage Upsampling (MSU) decoder architecture.** An input image (3 × H × W) is encoded by a frozen DINOv3 ViT backbone to produce a low-resolution spatial feature map (1280 x H/16 x W/16). The feature map is projected to 256 channels and progressively reconstructed to full resolution through a series of decoder blocks. Each block consists of a 3x3 convolution (followed by normalisation and SiLU activation) and a x2 bilinear upsampling operation. Finally, a 1x1 convolution maps the high-resolution features (64 x H x

W) to per-pixel class logits (N classes x H x W). All tensor shapes are shown in channel–height–width format.

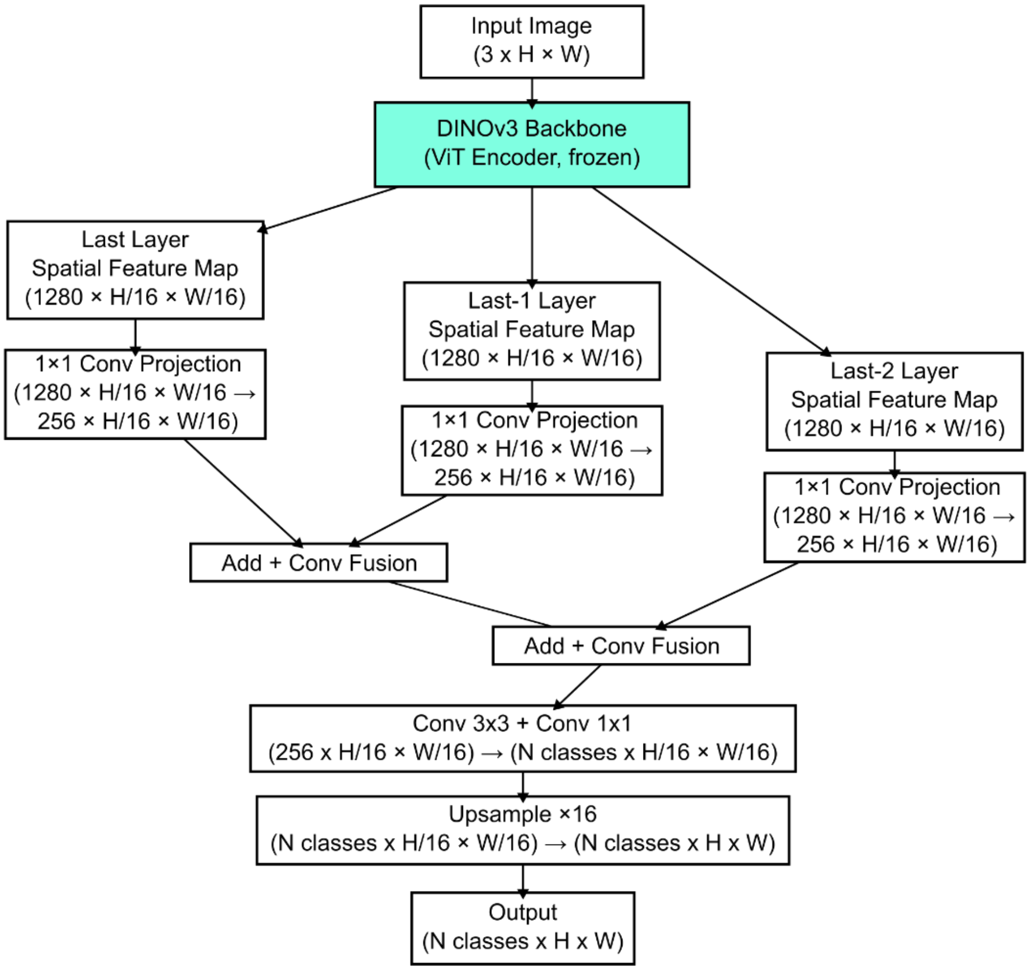


**Figure S4. Multi-layer Fusion (MLF) decoder architecture.** An input image (3 x H x W) is encoded by a frozen DINOv3 ViT backbone. Due to the isotropic nature of ViT, the extracted feature maps from the last n transformer layers all share the same spatial resolution (C × H/16 × W/16). These features are projected to 256 channels via 1x1 convolutions. The decoder performs a top-down fusion: starting from the deepest layer (Last Layer), features are progressively fused with shallower layers (e.g., Last-1, Last-2) via element-wise addition and convolution blocks. Finally, the fused feature map is projected to class logits and upsampled by x16 to generate the full-resolution segmentation masks (N classes x H x W). (Diagram illustrates n=3 for clarity, but we used the last four layers in our experiments*)*.

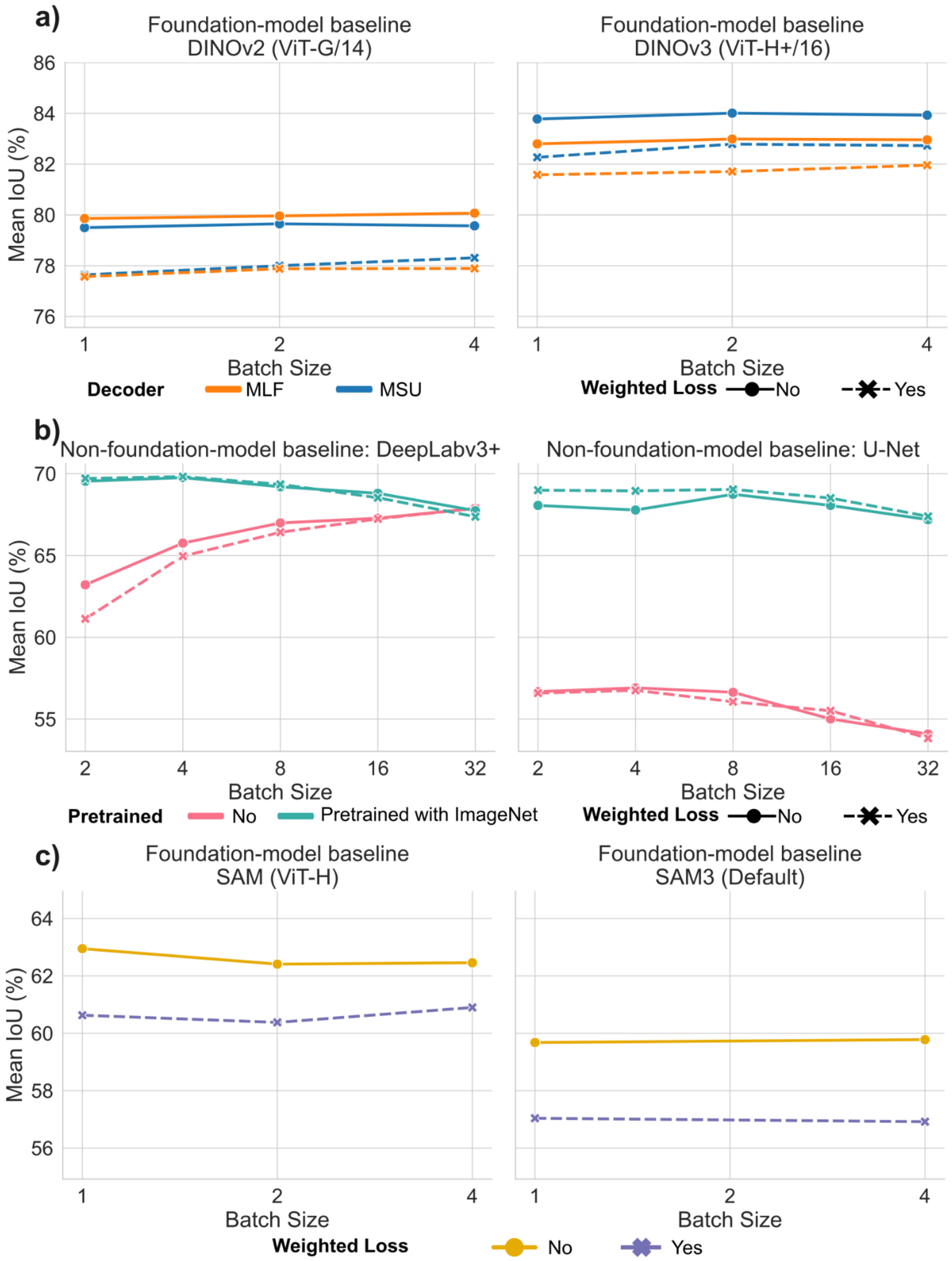


**Figure S5. Performance evaluation across segmentation architectures and training configurations.** (a) Foundation model baselines comparing DINOv2 (ViT-G/14) and DINOv3

(ViT-H+/16) encoders paired with Multi-Scale U-Net (MSU) versus Multi-Layer Fusion (MLF) decoders across batch sizes (N=1 to 4). (b) Classical fully supervised models (DeepLabv3+ and U-Net) were evaluated across batch sizes (N=2 to 32). Colours distinguish between random initialisation and ImageNet pretraining. (c) Performance of models using the Segment Anything Model (SAM) and SAM3 as encoder and MSU as decoder. Note: The data point for SAM3 at batch size 2 is omitted due to training instability (CUDA errors) consistently preventing convergence.

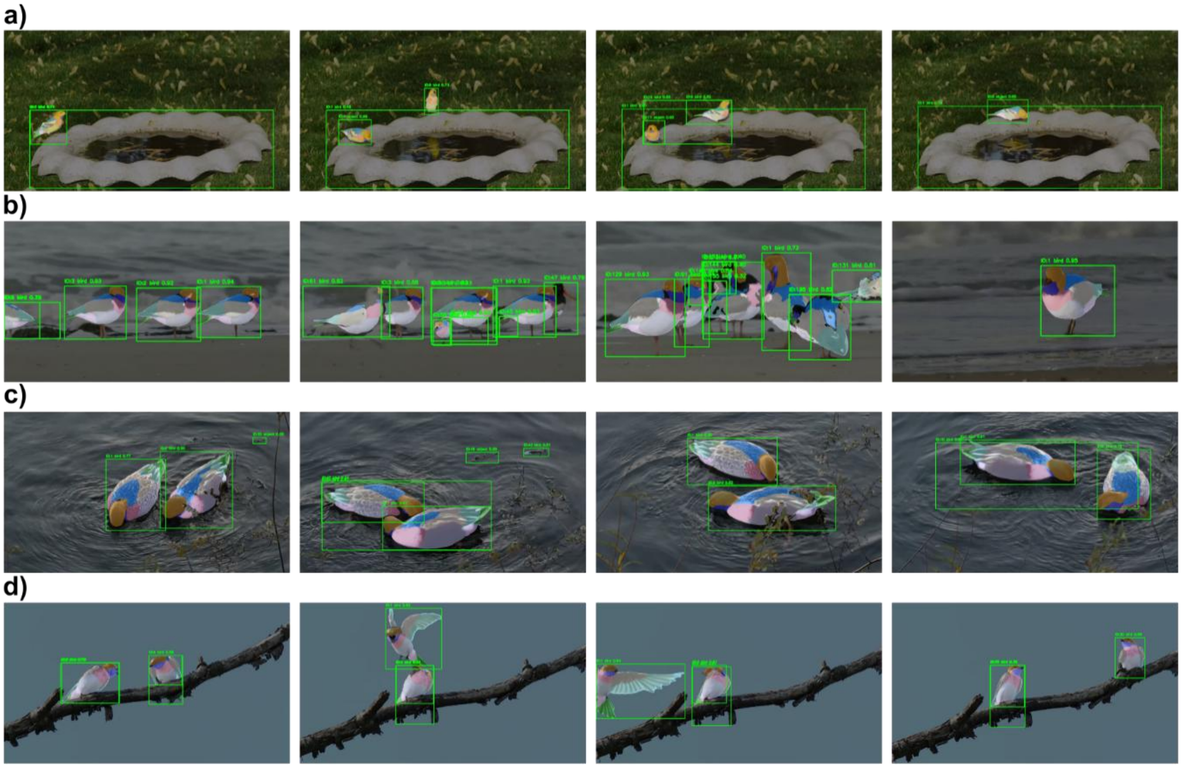


**Figure S6. Example results of our model applied to videos.** (a) Small targets: American Goldfinches (*Spinus tristis*) on a birdbath. (b) Density & Diversity: A flock of Eurasian Oystercatchers (*Haematopus ostralegus*) on a shoreline. This example highlights performance in crowded scenes with high visual diversity, including adult and juvenile, as well as a distinct species (a Black-headed Gull, *Chroicocephalus ridibundus*) visible in the second frame. (c) Occlusion handling: Northern Shovelers (*Spatula clypeata*) foraging in a river. The model successfully segments the plumage patches while correctly ignoring foreground occlusions (e.g., leaves covering the bird's body). (d) Pose variation: Movements of Tree Swallows (*Tachycineta bicolor*) during mating. The model maintains consistent segmentation masks across pose variations. Note: Green bounding boxes are raw outputs from the detection model. Image Credits & Licensing: Row (a) is from a video by J. M. Pearson

(CC0); Row (b) is adapted from a video by Natuur Digitaal (Marc Plomp) and Stichting Natuurbeelden (CC BY-SA 3.0 NL); Rows (c) and (d) are adapted from videos by Rhododendrites (CC BY-SA 4.0).
